# Serial dependence generalizes across the senses

**DOI:** 10.64898/2026.03.16.712008

**Authors:** Michele Fornaciai, Irene Togoli, Samuel Binisti, Olivier Collignon

## Abstract

Our perception often shows systematic biases revealing how the brain’s internal representation diverges from the physical world. In particular, what happened in the recent past can systematically attract current percepts, making them appear more similar to previous stimuli. Does this phenomenon reflect a low-level mechanism encapsulated within each sensory modality, or instead a higher-order post-perceptual decisional mechanism? Here we investigate the existence, mechanisms, and neural signature of serial dependence across vision and audition for numerosity perception. In a series of three complementary experiments, we show that serial dependence can transfer across modalities and is modulated by cross-modal attention. Electroencephalographic recordings further reveal that both uni-modal and cross-modal effects similarly emerge during the perceptual processing of the stimuli. Our results thus challenge both current mainstream accounts of serial dependence based on low-level and high-level computations, critically pointing to a central role of a functional, mid-level multisensory network in generating this phenomenon.

## Introduction

Our perception does not linearly reflect the external environment but instead shows systematic distortions that reveal the internal computations shaping our experience. For instance, our conscious perception is not composed of a series of isolated snapshots of the external world. Instead, each percept formed by our brain draws contributions from the spatial and temporal context that objects and scenes are embedded in. In particular, what we perceive in the present is systematically affected by what we perceived in the recent past. This leads to biases whereby a current percept appears to be more similar to a previous one than it actually is – a phenomenon known as *serial dependence*.

In vision, serial dependence has been demonstrated in virtually every domain, starting from fundamental features like orientation [1], color [2], motion [3], and numerosity [4,5], to more complex attributes like faces [6]. Serial dependence research in audition is more limited, but the effect has been demonstrated in domains like rate [7] and time [8] perception.

The ubiquity of serial dependence effects suggests that it reflects a fundamental and general brain computation; yet, the nature of this phenomenon remains elusive. On the one hand, serial dependence has been interpreted as reflecting a low-level mechanism supporting perceptual stability and continuity by smoothing out noise from sensory signals [1,9–11]. In striking contrast, serial dependence has been alternatively framed as occurring at a decisional, post-perceptual stage [12,13]. The nature of serial dependence effects thus remains an intensely debated and polarized topic, divided between low-level and high-level accounts. To resolve this debate and understand the nature and mechanistic origin of serial dependence, it is essential to understand the brain processing stages and brain networks involved in generating this bias. Does serial dependence operate within low-level networks involved with maintaining the stability of each sensory modality[8]? Or does it recruit a more widespread functional network, processing different but potentially redundant aspects of multi-sensory stimuli? Does it operate beyond perception entirely, involving abstract decisional representations [13]? Answering these questions is particularly important as it would allow to gain insights into the level at which this effect originates and operates, the computations supporting it, and the role that serial dependence plays in the complex hierarchy of brain processing. That is, it would allow to better understand the very nature of this phenomenon.

An appealing way to address this problem is to investigate the existence of cross-modal (e.g., audio-visual) serial dependence effects, as transferring information in time across the senses would evidence that those effects arise beyond the simple low-level features of the stimuli, and beyond the mechanisms dedicated to the stability of each given modality. According to a purely low-level sensory account, serial dependence should operate strictly within each sensory modality, not transferring across the senses. From the perspective of a high-level decisional account, instead, serial dependence is expected to transfer across sensory modalities, as it would operate on abstract decisional representations [14] detached from the modality they originate from. A third, middle-ground possibility is a stability mechanism operating at mid-level perceptual stages – that is, not on individual sensory features, but on integrated object or event representations including signals from multiple sensory modalities. In this case, cross-modal effects should emerge during the perceptual processing of a stimulus, before decision.

Results so far, although mixed, seem to point to the existence of modality-specific stability mechanisms, thus supporting the idea of serial dependence being encapsulated within low-level sensory networks that do not interact with each other [15,16]. Failure to find cross-modal effects in previous studies might however depend on methodological pitfalls, like for instance using stimuli that are too different from each other [15]. The existence of serial dependence effects transferring across modalities thus remains an open question. Addressing this possibility and the mechanisms involved represents a unique opportunity to better understand the brain processing level at which serial dependence operates, and whether this phenomenon reflects purely uni-sensory perceptual stability and continuity processes, or a higher-order mechanism.

In this study, we performed three complementary experiments investigating the existence, as well as the mechanisms and neural signature of serial dependence of numerosity information, within and across vision and audition. By combining psychophysics and electroencephalography, we aim not only at addressing cross-modal serial dependence at the behavioral level, but also at understanding the brain processing level at which this phenomenon arises. Our study therefore represents the most comprehensive investigation of uni– and cross-modal serial dependence of numerosity information, how attention might modulate those effects, and how they are represented in the human brain.

## Results

### Experiment 1

In Exp. 1 we used a numerosity discrimination task and modulated the task-relevance of different sensory modalities in two task conditions. In each trial, we presented a sequence of three stimuli (streams of brief visual flashes or auditory tones): a task-irrelevant inducer (7 or 20 events), followed by a constant reference (12 events) and a variable probe (6-24 events). Participants compared the reference and the probe and reported which one of the two was more numerous. In the uni-sensory task condition, participants exclusively judged visual reference and probe stimuli, while serial dependence was induced by either visual (within-modal effect) or auditory (cross-modal effect) inducer stimuli. In the multi-sensory task condition, instead, participants always compared and judged both visual and auditory stimuli. These two conditions were designed to engage either sensory modality-specific attention, or cross-modal attention. A depiction of the paradigm is shown in Fig. 1A, while the results are shown in Fig. 2.

**FIGURE 1.**
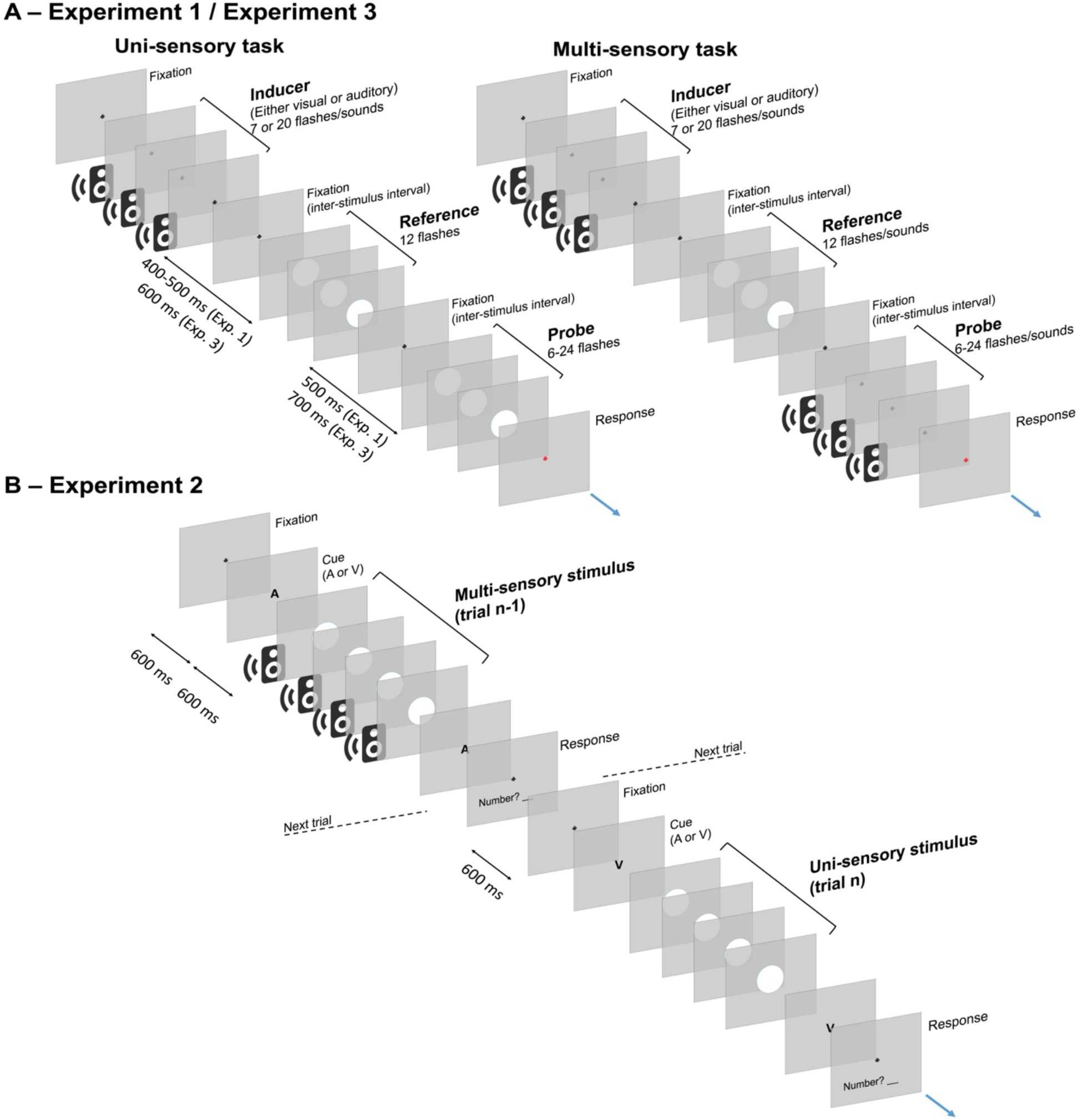
Experimental procedure. (A) General procedure used in Exp. 1 and Exp. 3. The experiment involved a discrimination task of sequential numerosities, and was divided into two different task conditions aimed at engaging modality-specific or cross-modal attention. In the “uni-sensory” task (left part of panel A), participants exclusively compared visual stimuli. The stimulus sequence involved a task-irrelevant inducer stimulus, which could be either a steam of brief visual flashes or a stream of brief sounds (numerosity = 7 or 20 events), followed by a constant reference (numerosity = 12) and a variable probe (numerosity = 6-24). Both the reference and the probe were always streams of visual flashes. The participants were instructed to report which stream (reference or probe) seemed more numerous. In the “multi-sensory” task condition (right part of panel A), the procedure was the same, but the reference and probe were always presented in different sensory modalities (a visual reference and an auditory probe, or vice versa). In both task conditions, the serial dependence effect was measured as the influence of the inducer on the perceived numerosity of the reference. (B) Procedure used in Exp. 2. The procedure in this case involved a numerosity estimation task. The trial sequence was structured in pairs of trials, in which we alternated multi-sensory (both auditory and visual sequences presented simultaneously) and uni-sensory stimuli. The components of the multisensory stimulus were modulated independently so that they could be congruent (both a low [4-9 events] or a high [16-26 events] numerosity) or incongruent (one low and the other high). A cue was presented 600 ms before the stimulus to inform the participants about which component they should attend and judge in each trial. Participants were instructed to report an estimate of the numerosity of the cued component of the multi-sensory stimulus, by typing a number using the keyboard. The uni-sensory stimulus had a numerosity drawn from an intermediate range compared to the multi-sensory stimulus (10-15 events). For consistency, a cue was presented also in this case before the stimulus. The task was the same as with the multi-sensory stimulus. The serial dependence effect in this case was computed considering the multi-sensory stimulus as the “previous” stimulus (trial n-1) and the uni-sensory stimulus as the “current” stimulus (trial n). Stimuli are not depicted in scale.

**FIGURE 2.**
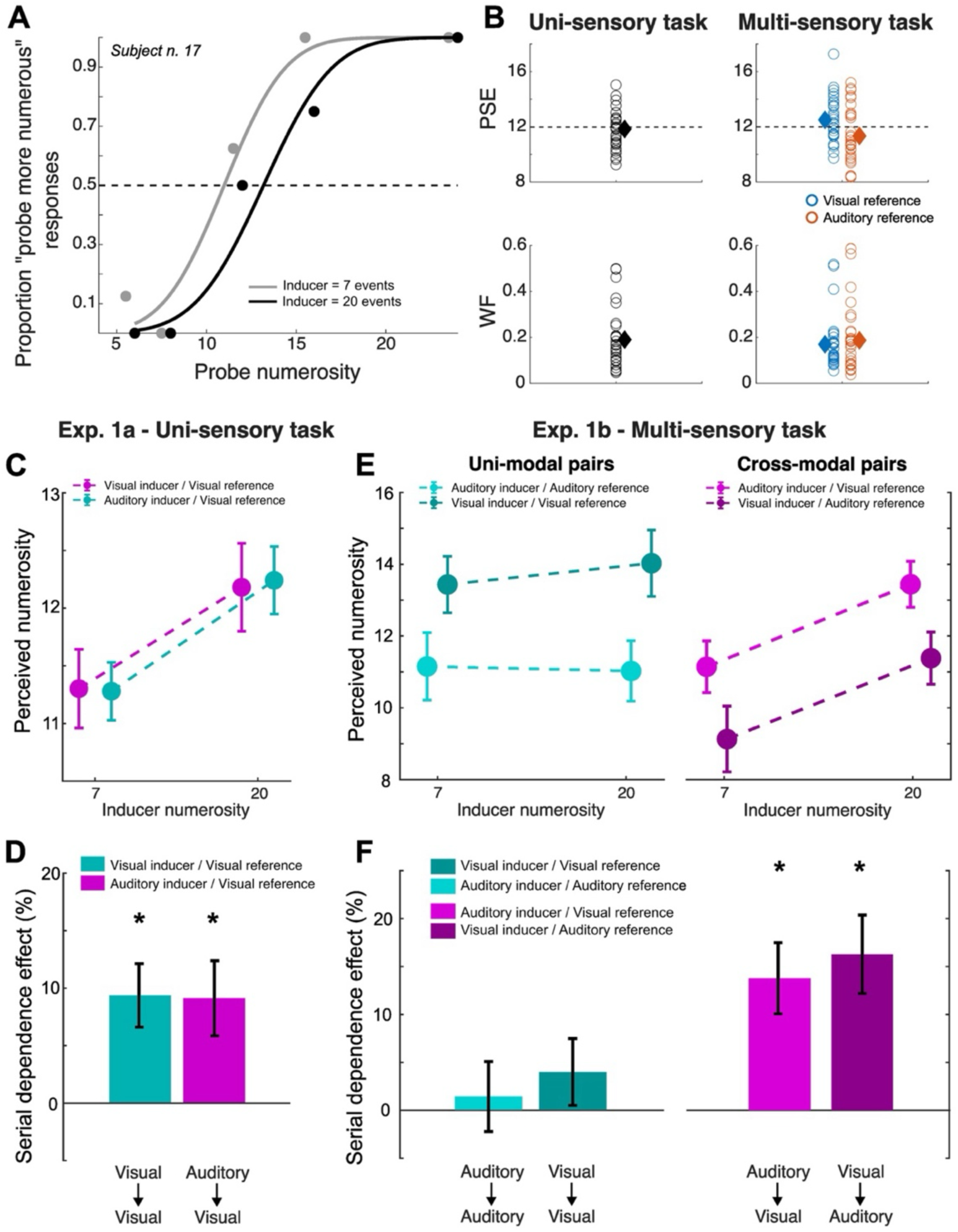
Results of Experiment 1. (A) Psychometric curves from a representative subject, computed in the uni-sensory condition of Exp. 1, and showing the effect of a visual inducer (within-modal effect). The dashed line indicates chance level (0.5). (B) General measures of performance in the two task conditions of Exp. 1. The two upper panels show the distribution (circles) and average (diamond) PSE in the uni-sensory and multi-sensory task (separately for the visual and auditory reference) condition. The dashed lines indicate the veridical numerosity of the reference (12). The two bottom panels show the distribution and average Weber’s fraction (WF). (C) Average PSEs as a function of the inducer numerosity, separately for the visual and auditory inducer in the uni-sensory task condition. (D) Average serial dependence effect indexes showing the effect of the visual and auditory inducer in the uni-sensory task condition. (E) Average PSEs as a function of inducer numerosity, corresponding to different combinations of inducer and reference modality in the multi-sensory task condition. (F) Average serial dependence effect indexes as a function of different combinations of inducer and reference modality, in the multi-sensory task condition. Error bars are SEM. * statistically significant effect.

Serial dependence in this paradigm was assessed as a shift in the perceived numerosity of the reference (point of subjective equality; PSE) according to the numerosity of the preceding inducer. Namely, with a low-numerosity inducer, we expected a relative underestimation of the reference numerosity, and a relative overestimation with a high-numerosity inducer. An example of the psychometric curves plotted to compute the PSE is shown in Fig. 2A. The psychometric curves of this representative participant indeed show a relative shift suggesting an attractive effect of the inducer on the perceived numerosity of the reference.

First, we assessed the general performance in the task (Fig. 2B). In terms of accuracy (reference perceived numerosity, PSE), in the uni-sensory task participants were on average very close to the veridical numerosity of the reference (PSE ± SD = 11.75 ± 1.29). In the multisensory task, we observed a slight overestimation of the visual reference (12.50 ± 1.65), and a slight underestimation of the auditory reference (11.34 ± 2.04). This is in line with previous results comparing visual and auditory numerosity perception [17]. The accuracy relative to the visual reference was significantly different from the uni-sensory task condition (paired t-test corrected with FDR; t(29) = –2.54, FDR adjusted-p = 0.032, Cohen’s d = 0.50), while the accuracy for the auditory reference was not (t(29) = 1.39, adj-p = 0.18). In terms of precision (Weber’s fraction, WF), we observed the best precision in the uni-sensory task (0.15 ± 0.08), while a slightly lower precision in the multi-sensory task (0.17 ± 0.11 and 0.19 ± 0.14, respectively for the visual and auditory reference), possibly reflecting the more demanding task of attending and comparing stimuli in different modalities. However, the difference was not statistically significant (p = 0.25 and 0.09, respectively).

We then assessed the participants’ perceived numerosity as a function of the preceding inducer. In the uni-sensory task condition, average PSEs show a marked difference as a function of the inducer, with a relative under and overestimation suggesting an attractive serial dependence effect (Fig. 2C). The difference in PSE appears very similar irrespective of the sensory modality of the inducer. The serial dependence effect indexes plotted in Fig. 2D better show the magnitude of serial dependence (i.e., normalized difference between the PSE obtained with high vs. low inducers, turned into percentage; ((PSE_high_ – PSE_low_)/PSE_low_) x 100). The visual and auditory inducer yielded a similar serial dependence effect (± SD) of 9.37% ± 15.12% and 9.12% ± 17.89%, respectively. Both effects are statistically significant (one-sample t-tests; t(29) = 3.39, adj-p = 0.004, d = 0.60; t(29) = 2.79, adj-p = 0.009, d = 0.50), with no difference between them (t(29) = – 0.09, p = 0.93). In the multi-sensory task condition, in terms of PSEs (Fig. 2E), we observed a different pattern of results depending on the modality of the inducer and reference stimulus. First, in line with what shown in Fig. 2B, visual reference stimuli were slightly overestimated compared to the veridical numerosity, while auditory stimuli were slightly underestimated. More interestingly, PSEs corresponding to uni-modal pairs of inducer and reference (within-modal effects; visual-visual, auditory-auditory) appear to be mostly flat, suggesting little serial dependence effect. This is indeed better shown by the average serial dependence indexes plotted in Fig. 2F. While cross-modal pairs of inducer and reference (auditory-visual, visual-auditory) showed robust and significant effects (11.47% ± 16.93%, (t(29) = 3.71, adj-p = 0.002, d = 0.66; 13.55% ± 18.65%, (t(29) = 3.98, adj-p = 0.002, d = 0.71), uni-modal pairs (visual-visual, auditory-auditory) showed weak, non-significant effects (1.15% ± 16.63%, (t(29) = 0.38, adj-p = 0.71; 3.29% ± 15.91%, (t(29) = 1.13, adj-p = 0.27). To better assess the difference between within-modal and cross-modal effects, we used a linear mixed-effect (LME) model, entering the serial dependence effect as the dependent variable, the modality of inducer and reference as the fixed effects, and the subjects as the random effect. The LME showed a significant interaction between the modality of inducer and reference (R^2^ = 0.09, b = –20.58, t = –3.36, p = 0.001), suggesting that only the combination of different modalities of past and present stimulus yield significant effects. Comparing the average cross-modal effect with the average within-modal effect (paired t-test) indeed showed a significant difference (t(29) = 3.16, p = 0.004, d = 0.83). In the multi-sensory task condition, thus, only cross-modal serial dependence effects emerged.

### Experiment 2

The results of Exp. 1 suggest that serial dependence in vision and audition is modulated by attention driven by the task. Namely, while both within-modal and cross-modal serial dependence effects emerge in a simple uni-sensory (visual) task, within-modal effects appear to be suppressed in a cross-modal (audio-visual) task.

In Exp. 2 we further assessed whether serial dependence within and across modalities could be modulated by attention in a trial-by-trial fashion, by means of attentional cueing. Namely, we alternated multi-sensory audio-visual stimuli and uni-sensory stimuli (either visual and auditory), and assessed the serial effect induced by each component of the multi-sensory stimulus. Crucially, before presenting the multi-sensory stimulus, we cued the component that participants had to attend and judge. See Fig. 1B for a depiction of the paradigm used in Exp. 2, while the results are shown in Fig. 3.

**FIGURE 3.**
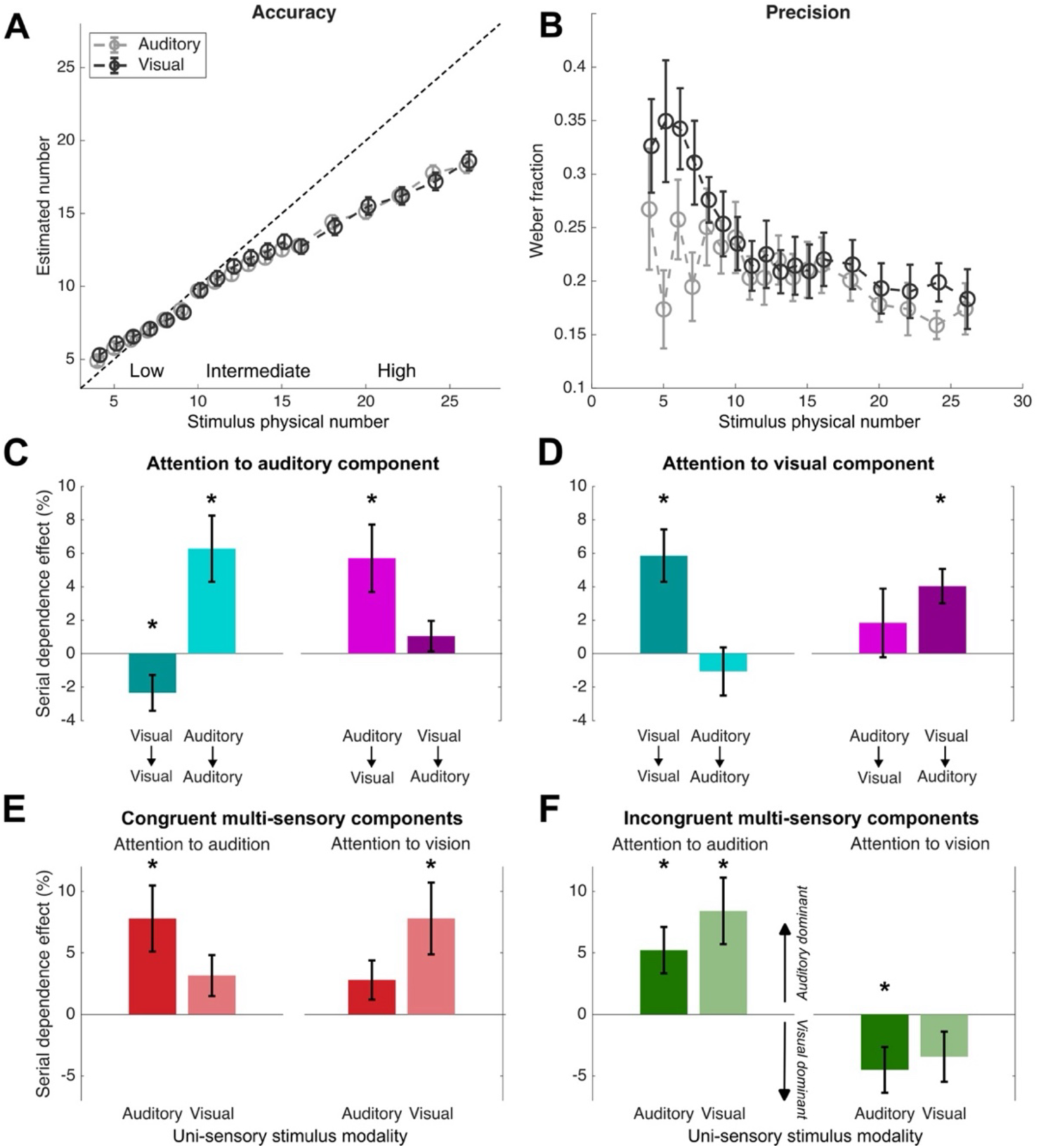
Results of Experiment 2. (A) General estimation performance in terms of accuracy (mean estimated numerosity). (B) Precision of numerical estimates across the ranges (Weber’s fraction, WF). (C) Serial dependence effects observed when attention was directed at the auditory component of the multi-sensory stimulus, as a function of the different components of the multi-sensory stimulus and modality of the uni-sensory stimulus. (D) Serial dependence effects observed when attention was directed at the visual component of the multi-sensory stimulus. (E) Effects observed when the two components of the multi-sensory stimulus were congruent (both low vs. both high numerosity), as a function of the attended modality and the modality of the subsequent uni-sensory stimulus. (F) Effects observed when the two components of the multi-sensory stimulus were incongruent (one low and the other high numerosity), as a function of the cued modality and the modality of the uni-sensory stimulus. The effects in this case were computed so that positive values indicate a dominant effect from the numerosity of the auditory component, while negative values indicate a dominant effect of the visual component. Error bars are SEM. * Statistically significant effect.

The range of numerosity of the stimuli used in this experiment was chosen so that each component of the multisensory stimulus could be either lower (4-9 events) or higher (16-26 events) than the subsequent uni-sensory stimulus (10-15 events). First, we thus assessed the accuracy of participants in estimating the numerosity of the different stimuli (i.e., the average estimated number at each level), as well as the precision of judgements (Weber’s fraction; WF = standard deviation / average judgment) at different numerosity levels and in different modalities. In terms of accuracy (Fig. 3A), we observed a typical pattern of estimation performance [4,18,19], with higher numerosities increasingly underestimated compared to the veridical numerosity. The results of a LME model (mean estimate ∼ numerosity level * modality + (1|subj)) showed only a main effect of numerosity (R^2^ = 0.89, b = 0.62, t = 25.48, p < 0.001), with no effect of stimulus modality (b = 0.33, t = 1.42, p = 0.16) or interaction (b = –0.01, t = –0.82, p = 0.41). In terms of precision (Fig. 3B), we observed higher WFs (i.e., lower precision) for visual stimuli in the low range, decreasing progressively towards higher numerosity levels. Auditory stimuli did not show such a sharp increase in WF at lower numbers, although with some variability. The results of a LME model (WF ∼ numerosity level * modality + (1|subj)) showed a significant interaction between numerosity and modality (R^2^ = 0.56, b = –0.004, t = –3.54, p < 0.001). To follow up this interaction, we performed a series of paired t-tests within each level of numerosity, comparing precision in vision versus audition. The results showed indeed a significantly better auditory precision at lower numerosities (5, 6, 7; p ≤ 0.031) and at 24 (p = 0.012). Numerosity estimation in the auditory modality thus showed better precision, although this was limited to a few levels of numerosity.

The serial dependence effect here was assessed as a difference in the estimated numerosity of the uni-sensory stimulus as a function of the numerosity of each component of the preceding multi-sensory stimulus. Fig. 3C-F shows the results in terms of serial dependence effect index, computed as the normalized difference in estimated numerosity when the preceding stimulus had a higher versus a lower numerosity. In addition, the effect was computed according to which component of the multi-sensory stimulus was cued, and the modality of the uni-sensory stimulus.

When attention was directed to the auditory component of the multi-sensory stimulus (Fig. 3C), only this component induced significant attractive serial dependence effects, irrespective of the modality of the subsequent stimulus. Namely, we observed significant effects from the auditory component on both subsequent auditory (6.27% ± 10.28%; t(26) = 3.17, adj-p = 0.016, d = 0.59) and visual (5.70% ± 10.52%; t(26) = 2.81, adj-p = 0.018, d = 0.52) stimuli. The visual (unattended) component of the stimulus, instead, had a small but significant repulsive (opposite) effect on a subsequent visual stimulus (−2.34% ± 5.53%; t(26) = –2.20, adj-p = 0.049, d = 0.41), and no significant effect on a subsequent auditory stimulus (1.05% ± 4.73%; t(26) = 1.15, adj-p = 0.26). A LME model on the effects as a function of the components of the multi-sensory stimulus and the modality of the uni-sensory stimulus (Effect ∼ Component modality*Uni. Modality + (1|Subj)) showed a significant main effect of the component of the multi-sensory stimulus (R^2^ = 0.18, b = 10.85, t = 2.26, p = 0.025), with no effect of the uni-sensory stimulus modality or interaction (p ≥ 0.20). An opposite pattern of effects was observed when attention was directed to the visual component of the preceding multi-sensory stimulus (Fig. 3D). That is, we observed significant effects from the visual component on both visual (5.86% ± 8.12%; t(26) = 3.75, adj-p = 0.002, d = 0.70) and auditory (4.03% ± 5.33%; t(26) = 3.93, adj-p = 0.002, d = 0.73) subsequent stimuli. No significant effect was induced by the auditory component on either an auditory (−1.06% ± 7.47%; t(26) = –0.73, adj-p = 0.48) or visual stimulus (1.84% ± 10.65%; t(26) = 0.89, adj-p = 0.48). A LME model however did not show any significant effect or interaction (R^2^ = 0.11, p ≥ 0.54), suggesting that in this case the pattern of effects is overall less clear. Serial dependence in the presence of a multi-sensory, audio-visual stimulus is thus governed by attention, and only the attended component of the stimulus seems to induce an attractive serial dependence effect. Moreover, the effect was roughly similar irrespective of whether it occurred within the same modality or across different modalities.

Additionally, we also addressed the possible effect of the congruence of the multi-sensory stimulus’ components. To do so, we sorted the trials according to whether the components were congruent (both low or high numerosity) or incongruent (one low and the other high numerosity), again as a function of the cued modality and the modality of the subsequent uni-sensory stimulus. Fig. 3E shows the effects induced by congruent multi-sensory stimuli. Interestingly, we observed a modulation of serial dependence, favoring within-modal effects. Namely, when the auditory modality was attended, the auditory component had a significant effect on a subsequent auditory stimulus (7.79% ± 13.96%; t(26) = 2.90, adj-p = 0.026, d = 0.54), but not on a subsequent visual stimulus (3.16% ± 8.66%; t(26) = 1.90, adj-p = 0.089). Vice versa, when the visual component was attended, a significant effect emerged only on a subsequent visual stimulus (7.80% ± 15.17%; t(26) = 2.67, adj-p = 0.026, d = 0.50), and not on an auditory stimulus (2.80% ± 8.26%; t(26) = 1.76, adj-p = 0.089). A LME model showed a significant interaction between the component of the multi-sensory stimulus considered to compute the effect, and the modality of the uni-sensory stimulus (R^2^ = 0.29, b = 9.63, t = 2.57, p = 0.011). On the other hand, when the components were incongruent (Fig. 3F), we observed a dominance of the cued component, which is indexed by positive serial dependence effect values when audition was attended, and negative values when vision was attended (see *Behavioral data analysis* for more information about how the index is computed). When the auditory component was cued, we found significant effects of the auditory component on both auditory (5.22% ± 9.81%; t(26) = 2.77, adj-p = 0.020, d = 0.52) and visual uni-sensory stimuli (8.41% ± 14.05%; t(26) = 3.11, adj-p = 0.016, d = 0.58). When the visual component was cued, we found a significant effect of this component on auditory uni-sensory stimuli (−4.50% ± 9.67%; t(26) = –2.42, adj-p = 0.031, d = 0.45), but not visual stimuli (−3.43% ± 10.59%; t(26) = –1.68, adj-p = 0.10). The pattern of effects may suggest stronger cross-modal effects in this case, that is, stronger effects on uni-sensory stimuli in a different modality compared to the cued modality of the multisensory stimulus. Paired t-tests however did not show any significant difference as a function of the modality of the uni-sensory stimulus (t(26) = 1.78, p = 0.086; t(26) = –1.51, p = 0.14; respectively for when the auditory and the visual component of the multi-sensory stimulus were cued).

### Experiment 3

After establishing the existence of cross-modal serial dependence and addressing the attentional mechanisms involved, we sought to understand the brain processing stage at which the effect emerges. This is indeed a key point to better understand the nature of this effect – that is, whether it involves perceptual or post-perceptual processing stages. In Exp. 3 we used the same psychophysical paradigm as Exp. 1 (which allows to more clearly distinguish between within-modal and cross-modal effects), but with the addition of electroencephalography (EEG) to address the neural signature of serial dependence.

In terms of behavioral results, the effects measured in Exp. 3 (Fig. 4) closely replicate what observed in Exp. 1. Namely, in the uni-sensory task (Exp. 3a; Fig. 4A) we observed robust serial dependence effects from both the visual (6.65% ± 14.34%) and auditory (10.19% ± 15.48%) inducers, again showing that in a simple uni-sensory visual task both within– and cross-modal effects occur. In both cases the effects were statistically significant (t(28) = 2.50, adj-p = 0.028, d = 0.45; t(28) = 3.55, adj-p = 0.003, d = 0.64), with no difference between them (t(28) = 0.90, adj-p = 0.37). In the multi-sensory task (Exp. 3b; Fig. 4B), on the other hand, we again observed robust cross-modal effects and reduced uni-modal effects. Namely, the effects observed considering cross-modal pairs of inducer and reference showed effect of 17.87% ± 26.79% (auditory-visual) and 14.53% ± 17.76% (visual-auditory). Both effects were statistically significant (t(28) = 4.41, adj-p = 0.002, d = 0.80; t(28) = 3.59, adj-p = 0.002, d = 0.65). Uni-modal pairs of inducer and reference instead showed effects of 6.42% ± 15.29% (auditory-auditory) and –1.19% ± 13.99% (visual-visual). In this case, the auditory within-modal effect was statistically significant (t(28) = 2.26, adj-p = 0.043, d = 0.40), while the visual effect not (t(28) = –0.45, adj-p = 0.65). Similarly to Exp. 1, however, a LME model including inducer and reference modality as fixed effects yielded a significant interaction between the two factors (R^2^ = 0.13, b = –27.17, t = –3.89, p < 0.001), suggesting that the combination of different modalities of the past and present stimulus resulted in stronger attractive serial dependence effects. This was confirmed with a paired t-test on the average cross-modal versus uni-modal effects, showing a significant difference (t(28) = 3.16, p = 0.004, d = 0.94). These results show again that attention driven by the task can modulate serial dependence, especially in a multi-sensory task.

**FIGURE 4.**
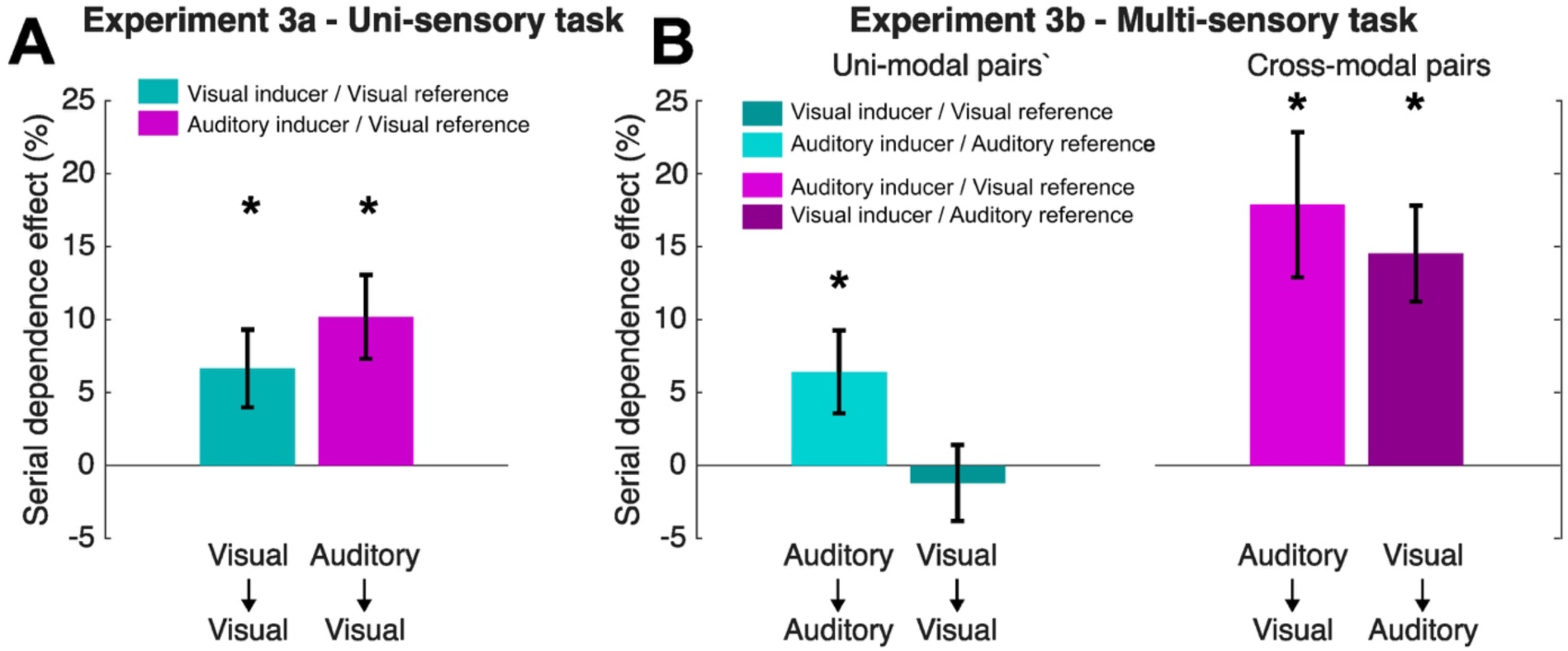
Behavioral results of Experiment 3. (A) Results of the uni-sensory task condition of Exp. 3, in terms of normalized serial dependence effect index. (B) Results of the multi-sensory task condition of Exp. 3. Error bars are SEM. * Statistically significant effect.

After assessing the pattern of behavioral effects, we went on and addressed the neural signature of serial dependence. The first step of our EEG approach was to select a group of channels of interest to perform further analyses. To do so, we examined the general responses to the visual reference stimulus in the uni-sensory task (Fig. 5A), and separately the visual and auditory references in the multi-sensory task (Fig. 6A-B). This procedure was chosen in line with our general hypothesis concerning the emergence of serial dependence in brain responses. Namely, if serial dependence affects the sensory processing of the reference stimulus, we wanted to select EEG channels that reflect the brain responses and processing of the reference, particularly at early stages (i.e., the first half of the reference presentation epoch). With this procedure, we selected a group of 12 channels showing peak responses to both visual and auditory references, including I1, Iz, I2, OI1, OIz, OI2, O1, Oz, O2, PO1, POz, PO2.

**FIGURE 5.**
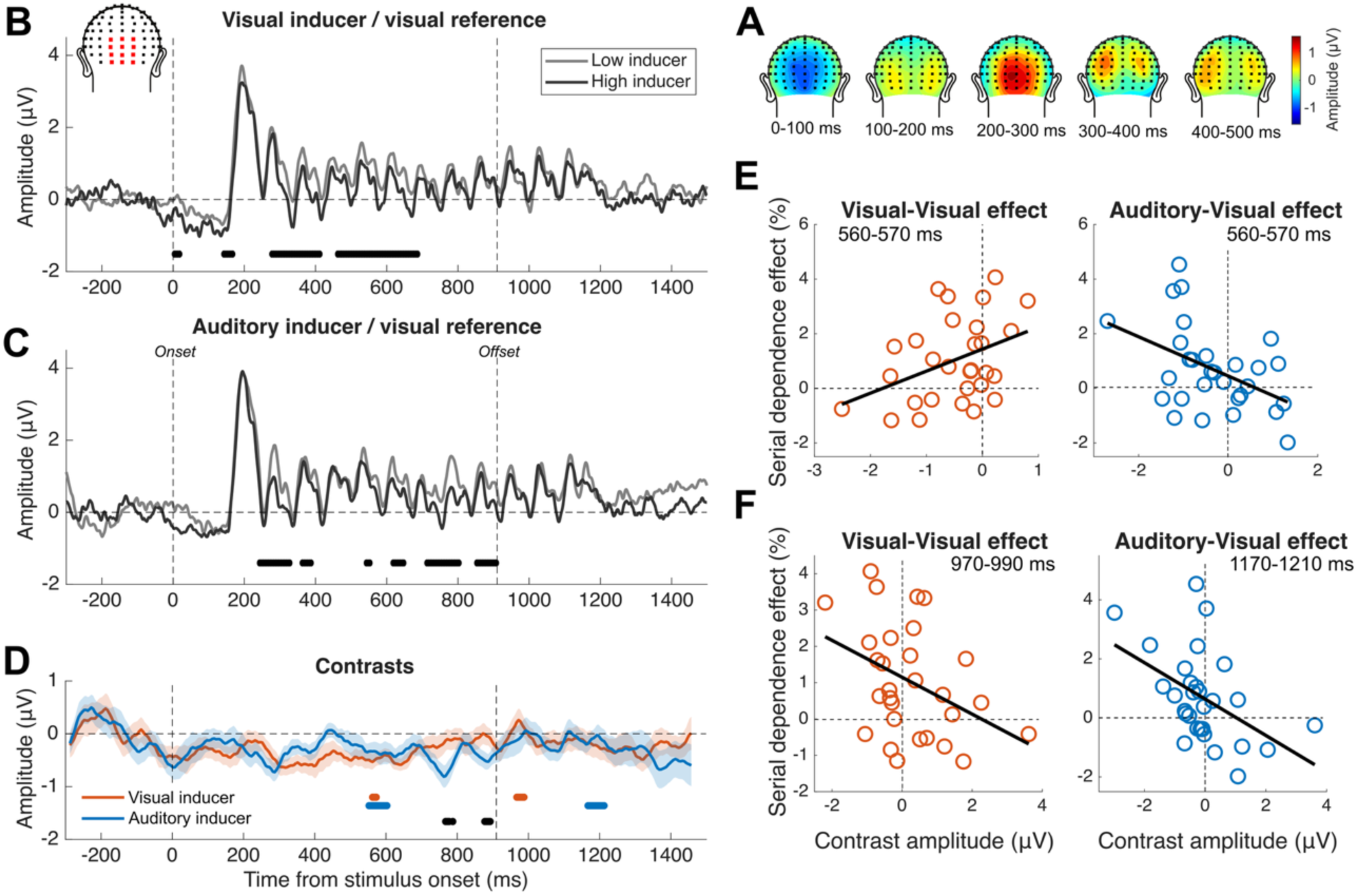
EEG results of the Uni-sensory task condition of Exp. 3 (Exp. 3a). (A) Topographic plots of activity evoked by the visual reference stimulus, in the first half of the reference presentation epoch. The topographic plots were used to select the channels of interest for further analysis. (B) Event-related potentials (ERPs) time-locked to the reference onset, corresponding to the combination of a visual inducer and a visual reference. The two waves correspond to the reference preceded by either a low-numerosity inducer (grey) or a high-numerosity inducer (black). The vertical dashed lines indicate the onset (0 ms) and offset (910 ms) of the reference. The horizontal dashed line indicates the zero in the amplitude scale. The inset in the top left corner shows the channel selected (I1, Iz, I2, OI1, OIz, OI2, O1, Oz, O2, PO1, POz, PO2). The waves plotted are the average of the selected channels. The black thick lines at the bottom of the plot show the significant latency windows observed with paired t-tests, showing a significant difference in ERP amplitude as a function of the inducer numerosity. (C) ERPs corresponding to the combination of auditory inducer and visual reference, divided according to the numerosity of the inducer. (D) Contrast waves relative to the effect of the visual (red) and auditory (blue) inducer, reflecting the difference (subtraction) of amplitude value of the pair of ERPs plotted in panel B and C, respectively. The black thick lines at the bottom of the plot mark the latency windows showing a significant difference between the two contrasts (paired t-tests). In all panels, the lines reporting the significance of tests span from the center of the first to the center of the last 50-ms window in which the tests were performed. The colored thick lines mark the latency windows showing a significant relationship between the contrast amplitude and the behavioral effect (difference in PSE as a function of the inducer numerosity). The shaded area represents the SEM. (E) Correlation between the contrast amplitude and behavioral serial dependence effect for the visual (left) and auditory (right) inducer, in a common latency window before the reference offset (560-570 ms). (E) Correlation between the contrast amplitude and behavioral serial dependence effect for the visual (left) and auditory (right) inducer, in two separate latency windows after the reference offset (970-990 ms and 1170-1210 ms, respectively).

**FIGURE 6.**
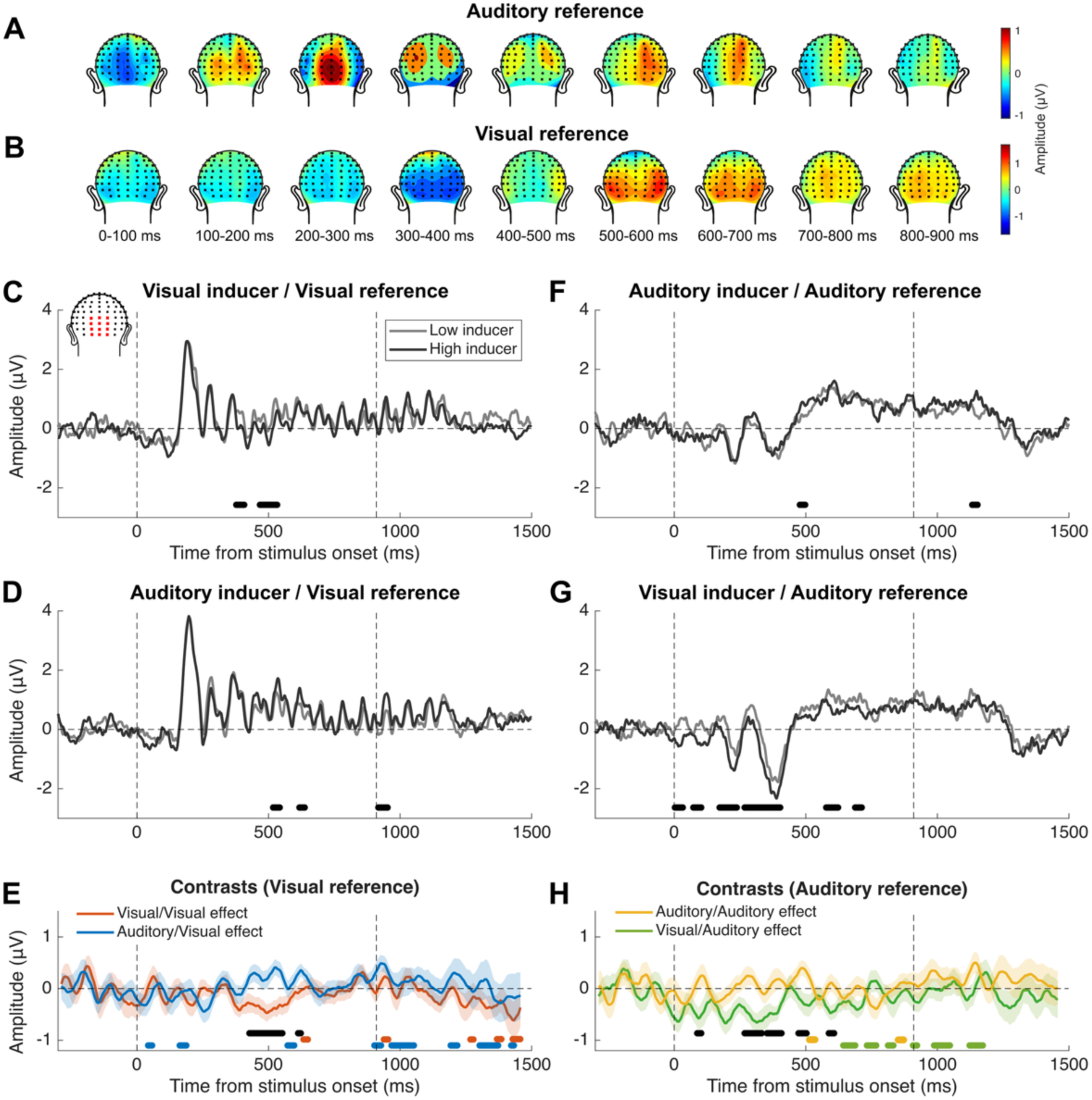
EEG results of the multi-sensory task condition of Exp. 3 (Exp. 3b). (A) Topographic plots of activity evoked the visual reference stimulus. (B) Topographic plots of activity evoked the auditory reference stimulus. (C) Event-related potentials (ERPs) time-locked to the reference onset, corresponding to the combination of a visual inducer and a visual reference. The two waves correspond to the reference preceded by either a low-numerosity inducer (grey) or a high-numerosity inducer (black). The vertical dashed lines indicate the onset (0 ms) and offset (910 ms) of the reference. The horizontal dashed line indicates the zero in the amplitude scale. The inset in the top left corner shows the channel selected. The waves plotted are the average of the selected channels. The black thick lines at the bottom of the plot show the significant latency windows observed with paired t-tests, indexing a significant difference in ERP amplitude as a function of the inducer numerosity. (D) ERPs corresponding to the combination of auditory inducer and visual reference, divided according to the numerosity of the inducer. (E) Contrast waves relative to the visual-visual (red) and auditory-visual (blue) combination of inducer and reference, reflecting the difference (subtraction) of amplitude values of the pair of ERPs plotted in panel C and D, respectively. The black thick lines at the bottom of the plot mark the latency windows showing a significant difference between the two contrasts (paired t-tests). The colored thick lines mark the latency windows showing a significant relationship between the contrast amplitude and the behavioral effect (difference in PSE as a function of the inducer numerosity). The shaded area represents the SEM. (F) ERPs corresponding to the combination of auditory inducer and auditory reference. (G) ERPs corresponding to the combination of visual inducer and auditory reference. (H) Contrast waves relative to the auditory-auditory (yellow) and visual-auditory (green) combination of inducer and reference, reflecting the difference (subtraction) of amplitude value of the pair of ERPs plotted in panel F and G, respectively.

First, in the uni-sensory task condition (Exp. 3a), the topographic plots of activity evoked by the reference stimulus (Fig. 5A) showed early activity (100-300 ms) centered at posterior occipital and occipito-parietal channels, followed by more lateralized activity. In line with this, we thus selected a set of posterior occipital/occipito-parietal channels (top-left inset in Fig. 5B). Then, we addressed the difference in the activity evoked by the reference as a function of the numerosity of the preceding stimulus. The reference was indeed constant across trials, so that any difference in the evoked response is likely to reflect the preceding stimulus. The significance of the effects was evaluated using a series of paired t-tests across a series of small time windows, in a sliding-window fashion (50 ms, step = 5 ms), controlled for multiple comparisons using a cluster-based non-parametric approach. The same sliding window testing and control for multiple comparisons was used for all other tests in the EEG analysis. First, the ERP plots (Fig. 5B-C) show a clear tracking of the numerosity of the reference, with individual peaks corresponding to each flash in the stream. Such peaks appear to be slightly delayed compared to the actual reference presentation timing, with some delay in the onset of the first peak and activity lingering after the offset. This however likely reflects the delay in activation of the visual cortex compared to the visual physical stimulation, due to conductance and processing times. In terms of effects, in the case of the within-modal effect (visual-visual; Fig. 5B) we observed four latency windows showing a significant difference in ERP amplitude as a function of the preceding inducer, starting from the onset of the reference (note that when reporting clusters of consecutive significant tests we report the center of the first and the last 50-ms window in which the tests were performed). Namely, we observed significant differences at 0-10 ms, 140-170 ms, 270-410 ms, and 460-690 ms (t = [min, max]; t(28) = [-4.31, –2.06], p ≤ 0.048). In all these latency windows, a preceding inducer with higher numerosity was associated with more negative (or less positive) amplitudes compared to a lower numerosity. In the case of the cross-modal effect (auditory-visual; Fig. 5C), we observed six significant latency windows, starting after 200 ms from the reference onset: 240-330 ms, 360-390 ms, 540-550 ms, 620-650 ms, 710-800 ms, and 850-910 ms (t(28) = [-4.13, –2.06], p ≤ 0.048). Again, higher inducer numerosity was associated with more negative ERP amplitudes. All these latency windows resulted significant in the cluster-based non-parametric tests used to control for multiple comparisons (all cluster p-values ≤ 0.002; see *Methods*).

To perform further analyses, we computed a measure of ERP contrast that reflects the difference in ERP amplitude as a function of the inducer numerosity (Fig. 5D). The contrast measures were first used to compare the visual and auditory inducer effects on the activity evoked by the reference. This showed two small latency windows before the offset of the reference, in which effects were significantly different: 770-780 ms, and 880-890 ms (t(28) = [-2.36, –2.05], p ≤ 0.048; both cluster p-values < 0.001). In both windows, the effect of the auditory inducer showed increased amplitude with negative polarity. Moreover, the contrasts were used to assess the relationship between the inducer effect at the neural level, and its effect at the behavioral level. To do so we computed the difference in PSE as a function of the inducer numerosity as an index of the behavioral effect, and used a LME model to assess if this effect can be predicted by the difference in ERP amplitude (i.e., the contrast). In the model, the behavioral effect was entered as the dependent variable, while the ERP contrast as the fixed effect, with the addition of the subjects as the random effect. The tests were performed throughout the reference presentation epoch in a series of small time windows (50 ms, step = 5 ms), in a sliding-window fashion. Interestingly, we observed a significant relationship between neural and behavioral measures of the effect of both the visual and auditory inducer in overlapping latency windows during the reference presentation epoch. Specifically, at 560-570 ms (R^2^ ≥ 0.62; b = [0.74, 0.81], t = [2.14, 2.28], p ≤ 0.043), and 550-600 ms (R^2^ ≥ 0.61; b = [-0.71, – 0.49], t = [-2.83, –2.09], p ≤ 0.030]), respectively (note that the results are reported indicating [min, max] of beta and t-values). The visual inducer showed an additional significant window at 970-990 ms (R^2^ ≥ 0.61; b = [-0.57, –0.51], t = [-2.37, –2.15], p ≤ 0.040), after the offset of the reference, while the auditory inducer at 1170-1210 ms (R^2^ ≥ 0.61; b = [-0.97, –0.50], t = [-3.50, –2.31], p ≤ 0.030). All these latency windows resulted significant in the cluster-based non-parametric tests (cluster p-values < 0.001). These results show that the extent to which the inducer modulates the brain responses to the reference is linked to the strength of the serial dependence effect measured behaviorally.

We further examined these latency windows individually by averaging the activity within them, and performing a correlation test with the behavioral effect. To do so, we chose a common early latency window (before stimulus offset) for the visual and auditory inducer (560-570 ms), and two different later windows (after stimulus offset; 970-990 ms and 1170-1210 ms, respectively for the visual and auditory inducer). At the early common window, the results confirmed a significant correlation between the contrast amplitude and the behavioral effect, but, interestingly, in two different directions. Indeed, for the visual inducer, the behavioral effect is positively associated with the contrast amplitude, increasing for more positive (or less negative) contrast amplitudes (r = 0.39, p = 0.036). For the auditory inducer, instead, the association is negative, with the effect increasing with increasingly more negative contrast amplitudes (r = –0.45, p = 0.014). At the two later latencies, we observed again significant correlations, both in the negative direction (r = –0.39, p = 0.035; r = –0.47, p = 0.009, respectively for the visual and auditory inducer).

The results from the uni-sensory task condition overall show a clear signature of the effect of the inducer in ERPs, and a relationship between such signature and the behavioral effect. The multi-sensory task condition offers the opportunity to further expand these findings with a more comprehensive set of combinations of sensory modalities, and to better understand the effect of attention. Similarly to the uni-sensory condition, we first plotted the topographic distribution of activity evoked by the reference. This was done separately for the auditory (Fig. 6A) and visual (Fig. 6B) reference stimuli. Both types of reference stimuli showed strong occipital and occipito-parietal activity, with both central and more lateralized responses. In line with the procedure used in the uni-sensory task, we again chose a set of central occipital/occipito-parietal channels, for both visual and auditory stimuli. The channels selected were the same as the uni-sensory task condition, and are shown in the top-left inset in Fig. 6C. All the plots and analyses were based on the average activity from these channels.

In terms of ERPs reflecting the effect of the inducer on the activity evoked by the reference, first we assessed the case of the visual reference, which was similar to the uni-modal task condition. Again, the activity evoked by the reference (Fig. 6C-D) showed a clear tracking of the numerosity of the stimulus, with individual peaks for each flash in the reference stream, slightly delayed compared to the actual presentation timing of the stimuli. Even if it was not effective in terms of behavioral effect, the visual inducer (Fig. 6C) yielded two latency windows showing a significant effect (380-440 ms and 470-530 ms; t(28) = [-2.98, –2.05], p ≤ 0.049). The auditory inducer (Fig. 6D) instead showed three significant latency windows, the first two close to the visual effect (520-540 ms and 620-640 ms; t(28) = [2.06, 2.78], p ≤ 0.048), and one later window around the reference offset (920-950 ms; t(28) = [2.12, 3.04], p ≤ 0.043). It is worth noting in this context that visual (within-modal) and auditory (cross-modal) effects on the visual reference showed opposite polarities. Namely, while the effect of the visual inducer was associated with more negative amplitudes for the higher numerosity, the effect of the auditory inducer was positive.

In terms of contrast amplitudes (Fig. 6E), we observed a large significant difference between the visual and auditory effects, at 430-550 ms (t(28) = [2.23, 4.93], p ≤ 0.038), which showed opposite polarities within this window. An additional small significant window was observed at 610-620 ms (t(28) = [2.16, 2.20], p ≤ 0.039). In terms of relationship between the neural (i.e., contrasts) and behavioral effects, both the visual and auditory inducers showed several significant latency windows. Namely, for the visual inducer, a significant relationship emerged at five latency windows, one before the reference offset (620-640 ms), and the rest at later latencies after the offset (940-950 ms, 1270-1280 ms, 1370-1380 ms, 1430-1460 ms; R^2^ ≤ 0.61; b = [-0.64, –0.26], t = [-3.29, –2.05], p ≤ 0.049). In all these windows, the relationship between neural and behavioral effects was negative, with the strength of serial dependence increasing for increasingly negative contrast amplitudes. For the auditory inducer, a significant relationship emerged at eight latency windows, starting very early right after the onset of the reference stimulus. The four early windows within the reference presentation epochs (40-60 ms, 160-190 ms, 570-600 ms, and 900-930 ms) showed a positive relationship between the contrast amplitude and the serial dependence effect (R^2^ ≥ 0.61; b = [0.46, 0.76], t = [2.07, 2.98], p ≤ 0.048). The later latency windows after the offset of the reference (970-1050 ms, 1190-1220 ms, 1300-1370 ms, and 1420-1460 ms) showed instead a negative relationship (R^2^ ≥ 0.61; b = [-0.56, –0.20], t = [-3.04, –2.08], p ≤ 0.044). All these latency windows resulted to be significant in the cluster-based non-parametric tests (cluster p-values < 0.001).

The activity evoked by the auditory reference (Fig. 6F-G) generally showed a different dynamic compared to the visual reference, and in particular it lacked the clear tracking of the reference events. Instead, it showed two main negative peaks at around 250 and 450 ms, followed by a sustained positive activity. The auditory inducer had little effects on the auditory reference, with only two small latency windows showing a significant difference as a function of the inducer numerosity (450-520 ms and 1100-1170 ms; t(28) = [2.22, 2.71], p ≤ 0.034). The visual inducer instead had a much more robust and extensive effect, with six significant latency windows starting right after the onset of the reference (0-55 ms, 60-120 ms, 140-235 ms, 240-420 ms, 550-640 ms, and 670-730 ms; t(28) = [-3.80, –2.09], p ≤ 0.045). Interestingly, similarly to the case of the visual reference, the ERP effect of auditory and visual inducers on the auditory reference showed different directions, but reversed compared to the visual reference. Namely, in this case, the auditory (within-modal) effect showed a positive polarity, while the visual (cross-modal) effect a negative polarity. All cluster p-values for these tests were < 0.001.

In terms of ERP contrasts, we observed again differences between the effect of the auditory and visual inducer, at five latency windows (60-120 ms, 230-350 ms, 355-430 ms, 440-520 ms, 560-630 ms; t(28) = [-3.70, –2.05], p ≤ 0.049). We also observed multiple latency windows showing a relationship between the neural and behavioral effect. For the auditory inducer, which had very limited behavioral effects, we observed only two small latency windows at 500-550 ms (R^2^ ≥ 0.62; b = [0.82, 0.91], t = [2.12, 2.38], p ≤ 0.042) and 830-900 ms (R^2^ ≥ 0.63; b = [-0.95, –0.87], t = [– 2.57, –2.15], p ≤ 0.040), showing relationships in different directions (i.e., positive and negative). For the visual inducer, instead, we observed six significant windows, with the first two at 620-710 ms and 700-790 ms showing a negative relationship between the contrast amplitude and the strength of serial dependence (R^2^ ≥ 0.61; b = [-1.72, –0.94], t = [-3.50, –2.09], p ≤ 0.045), and the other four (800-850 ms, 900-950 ms, 970-1070 ms, and 1100-1190 ms) showing a positive relationship (R^2^ ≥ 0.62; b = [0.66, 1.31], t = [2.08, 3.34], p ≤ 0.047). Overall, also in the multi-sensory task condition we observed a robust relationship between the neural effects of the inducer and the behavioral serial dependence effect, starting from relatively early latencies before the offset of the reference stimulus.

## Discussion

Our perceptual experience is not veridical but shows systematic distortions that reveal how the brain’s internal representations diverge from the physical world. For instance, perceiving stimuli in the present often shows attractive biases toward recent sensory experiences. The origin of such serial dependence effects remains debated [1,13]. Some have proposed that serial dependence reflects a stability mechanism [9,10] encapsulated in low-level sensory processes [15,16,20] while others suggest a high-level mechanism based on decision and/or memory [12,13]. Overall, by combining psychophysical and electroencephalographic experiments, we show the existence of cross-modal serial dependence effects, emerging at an intermediate perceptual stage of information processing, and gated by attention. Our results demonstrate that serial dependence effects can flexibly transfer across the senses based on task demands, revealing that current perception is influenced by experience embedded into functionally specific and interacting multisensory networks dedicated to solving a similar task, like numerosity perception. Yet, such an influence plays out during perceptual processing, suggesting that it alters our phenomenological experience of the stimuli rather than our decisions.

So far, only limited evidence has been provided concerning the existence of cross-modal serial dependence, and mostly based on properties that are intrinsically abstracted from the sensory modalities (e.g., effects induced by subjective ratings of emotional valence) [14,21]. Studies addressing the effect on perceptual features like numerosity have instead failed to find evidence for cross-modal effects [15,16], supporting the idea that this phenomenon remains encapsulated in sensory-specific brain networks. However, the use of sub-optimal stimuli (i.e., relying on different presentation formats) [15,22] or perceptual dimensions (i.e., time, which often shows weaker effects compared to other dimensions) [16,23] may explain the apparent absence of evidence. Clarifying the potential existence of cross-modal effects and their neural correlates is however pivotal to understand the nature and mechanisms of serial dependence. Indeed, the idea of a low-level stability mechanism [1,24,25] would predict effects that do not transfer across modalities. A high-level decisional mechanism would instead predict effects arising across modalities [13], although with them being driven by decisional representations rather than a genuine cross-modal transfer of perceptual information. A third, middle-ground possibility is the existence of cross-modal effects emerging from a functional mid-level network, supporting the stability of global, integrated perceptual representations rather than low-level features per se. Whether or not cross-modal serial dependence effects exist – and at which processing stage they potentially emerge – can thus provide important evidence for the nature and role of this phenomenon in perception and decision-making.

In this context, attention represents another important factor to consider due to its intrinsic link with multisensory integration [26]. Attention may indeed support or inhibit cross-modal effects, and can potentially provide additional insights into the mechanisms of serial dependence. Previous results show that serial dependence is strongly modulated by attention [1,5,27], making it an important component of this phenomenon. Thus, understanding how cross-modal attention gates serial dependence effects across the senses may provide additional evidence for the role of modality-specific and multisensory brain networks.

To address these issues, in a series of three complementary experiments we focused on the existence of cross-modal serial dependence effects in vision and audition, and the attentional mechanisms governing them. Our behavioral results first confirm that serial dependence occurs cross-modally in numerosity perception, showing that this effect is not limited to operating within a given sensory modality (see Fig. 2-4). Differently from previous studies [15], we show that stimuli with a similar spatio-temporal structure (i.e., fully sequential numerosities) presented in different modalities can affect each other attractively. This is in line with recent results showing that numerical brain representations align only when vision and audition share a sequential format [22]. A key to achieve multisensory integration of past and present stimuli seems thus to have stimuli in a similar presentation format, giving rise to aligned neural representations that can interact with each other.

Since our experience of the world is often multisensory, such temporal influence between auditory and visual numerosity may allow an efficient interaction between the senses and support fast and optimal representation of multisensory numerosity signals. Indeed, serial dependence likely reflects an adaptive process aiding the stability, consistency, and continuity of our experience in a noisy world [10]. These results thus point to the idea that serial dependence in perception does not operate at a low, featural level fully contained within a given modality, but stabilizes instead an integrated representation of multisensory events. Considering that information from multiple senses (and especially vision and audition) is often redundant, this cross-modal transfer suggests an efficient stability mechanism leveraging all available information concerning audio-visual events. Interestingly, recent results show that serial dependence does not improve behavioral performance, but tends to deteriorate it [28]. A stability mechanism however is not necessarily expected to improve behavior. A degraded performance might in fact represent the necessary trade-off to enable a continuous and consistent phenomenological experience in face of noise and random fluctuations in neuronal signals.

Our findings also show that the phenomenon can be modulated by task-driven attention (see Fig 2-4). Interestingly, it is not the cross-modal effect that is mostly impacted by attention, but the effect occurring within each sensory modality. Based on previous results concerning the effect of attention in visual serial dependence [27] and multi-sensory integration [29,30], one might expect modality-specific attention to suppress cross-modal effects. Instead, our results show that attending a single task-relevant modality does not prevent cross-modal effects. In fact, cross-modal (audio-visual) effects in our visual task were as strong as within-modal (visual-visual) effects. It is instead cross-modal attention that seems to strongly modulate serial dependence. Indeed, in the multi-sensory audio-visual task (Exp. 1b) we observed much reduced, almost completely suppressed, within-modal effects. This finding was also closely replicated in Exp. 3, suggesting that this is indeed a robust result. One possibility explaining this pattern of results is that in a simpler uni-sensory visual task either the auditory stimuli could implicitly capture attention leading to serial dependence, or the sensory systems could have more resources to maintain and integrate past information from different sources. In a more engaging multi-sensory task, the mechanisms responsible for serial dependence might instead become more selective, maintaining and integrating only “relevant” information – in this case, information from a different modality. This would occur via a stronger multisensory integration of consistent (i.e., similar) signals originating from different sources. Indeed, under conditions of cross-modal attention, the brain preferentially integrates complementary signals from different modalities since they have independent levels of noise, making the integration process more robust. This is supported by previous results showing a stronger integration of redundant multisensory cues rather than uni-sensory cues, leading to enhanced integrative gains [31].

In general, attentional and cognitive load has been shown to strengthen multisensory integration effects [32,33], but this does not explain the suppression of within-modal effects. Instead, we believe it more likely that such a modulation reflects an active gating of information to optimize integration, prioritizing different sources according to the context. In line with this idea, in a recent study from our group using multi-dimensional stimuli (i.e., dot arrays varied in numerosity, duration, and size) [23], we observed that only the task relevant stimulus dimension induces a serial dependence effect. Nevertheless, information from the other dimensions is still encoded in brain (EEG) responses, at least to some extent. This supports the idea that serial dependence is governed by an active gating mechanism selecting relevant past information for integration with current perceptual representations.

By modulating modality-specific attention in a trial-by-trial fashion (Exp. 2), we show that, on average, only the attended component (either visual or auditory) of a past multi-sensory stimulus induces a serial dependence effect. This effect emerges irrespective of the modality of the subsequent stimulus, in line with the results of Exp. 1a. However, when considering the congruency of the different components of the multi-sensory stimulus, a more complex picture emerges. Namely, with congruent auditory and visual components (both high or both low numerosity), we observed an enhanced selectivity for the sensory modality of the subsequent stimulus, favoring within-modal effects. In line with the interpretation of Exp. 1, this might be due to an increased selectivity in the use of past information in more engaging conditions. When the components are similar in numerosity and unfolds in parallel, indeed, they might more easily compete for resources, with the irrelevant modality capturing attentional resources. With different, incongruent numerosities, instead, the relevant modality is much more easily separable from the irrelevant one, due to the widely different ranges that we used. As the cue engaged modality-specific attention, this increased selectivity resulted in enhanced within-modality effects. If this interpretation holds true, it predicts that having a more engaging setup also in a uni-sensory task like Exp. 1a (for instance by introducing a secondary task), should increase the selectivity of the effect in favor of within-modality effects. This is an interesting possibility that should be tested in future studies.

Overall, our behavioral results demonstrate that serial dependence can transfer across sensory modalities, and is gated by attention depending on the task performed. Central to the aims of our study is to address the nature of this phenomenon. The existence of a cross-modal transfer rules out a purely low-level sensory effect encapsulated into modality-specific processing. However, is the effect therefore arising from abstract decisional representations? Or is it a perceptual effect occurring in a mid-level multisensory brain network? Our EEG results show the processing stage at which the serial dependence effect occurs. Specifically, our findings show a signature of the effect emerging at latencies even before the offset of the reference stimulus. This is particularly interesting because at the time the effect emerges from EEG signals, the stimulus sequence is still being displayed on the screen or played from the speakers. In fact, most of the effects are evident within the first half of the stimulus presentation, and in some cases even shortly after the onset of the sequence (e.g., see Fig. 5). This suggests that serial dependence affects the perceptual processing of the stimulus, rather than the decision based on it. At the time the effect emerges, a decision is indeed unlikely to occur, since a good portion of the stimulus still needs to unfold. A decision will only be possible once the full sequence has unfolded, or at least in the last part of the sequence (see for instance [34] for a signature of decision in duration perception). A likely candidate explaining the effect at the perceptual level is the accumulation of numerosity information over time, which has been shown to be affected by the spatio-temporal properties of the stimuli [35,36]. A difference in the rate of accumulation based on perceptual history can lead to differences in perceived numerosity, pushing the appearance of the current stimulus towards the previous one. This finding is also in line with previous studies showing a relatively early neural (EEG) signature of serial dependence [23,37,38], and with fMRI results showing the involvement of the primary visual cortex [39]. Additionally, the results also show a robust relationship between the neural effect (i.e., change in ERP response amplitude) and the behavioral effect. That is, the modulation of ERP amplitude shown in our results is significantly related to a change in the strength of the behavioral bias, and thus, effectively a change in the perceived magnitude of the stimuli. This is particularly important as it suggests that the brain processes occurring at such stages are indeed relevant for perception and behavior rather than being an epiphenomenon, and likely involved in determining the perceived numerosity of the reference sequence.

In terms of a broader framework of serial dependence effects, our results overall suggest that serial dependence operates at an intermediate perceptual stage, at least when it comes to perceptual dimensions such as numerosity [25,40,41]. For instance, the cross-modal effect may involve a direct exchange of signals across a functional multisensory brain network including visual and auditory cortices. In principle, the encoding and transmission of past information may additionally involve feedback from higher-level areas like the intraparietal sulcus [42–45], which would be in line with previous psychophysical results [25,40,41]. Irrespective from the potential involvement of feedback signals, the key aspect of our results is that the effect emerges at a stage consistent with the formation of perceptual representations rather than later decision-making processes. Finally, attention might modulate the coupling between the different sensory cortices within the multisensory network, governing the effect according to the current task demands – something similar to how attention modulates the coupling of sensory areas in multisensory integration [26,46].

To conclude, our results demonstrate that serial dependence can transfer across vision and audition in numerosity perception. We further demonstrate that effects within and across modalities are gated by attention. Finally, our EEG results show that the effect emerges during the perceptual processing of the current stimulus, possibly affecting processes like the accumulation of sequential numerosity information. Taken together, our results show that the perceptual systems are remarkably flexible in integrating past and present stimuli, leveraging different sources and gating information according to its relevancy and consistency with the current behavioral goals. Crucially, our results challenge the notion that serial dependence is encapsulated in low-level sensory stages of information processing, and instead suggests that current perceptual representations are contextualized by interacting multisensory networks involved in solving similar computations, like estimating numerosity.

## Methods

### Participants

A total of 91 participants took part in the study (72 females, age range 18-36 years). 32 participants were tested in Exp. 1, 27 participants in Exp. 2, and 32 participants in Exp. 3. All participants had normal or corrected-to-normal vision and audition, and reported no history of neurological, developmental, or psychiatric disorder. Each participant read and signed a written informed consent form before the start of the session. The study was approved by the ethics committee of Université Catholique de Louvain (protocol #2023-48), and was designed to be in line with the declaration of Helsinki. Two participants were excluded from data analysis in Exp. 1 due to insufficient performance (Weber’s fraction > 1). Three participants were excluded in Exp. 3, two because of insufficient performance, and one because of too noisy EEG data. The sample size tested in each experiment was computed a priori according to a power analysis based on the typical effect size of serial dependence in numerosity perception. Namely, we computed the average effect size (Cohen’s d) from multiple previous studies using similar numerosity discrimination or estimation paradigms [5,15,19,23,37,40,47], which resulted to be d = 0.71. A power analysis considering a two-tailed distribution and a power of 0.9 yielded a minimum sample size of 24 participants, which we slightly increased to be more conservative and to account for the possible exclusion of a few participants.

### Apparatus and stimuli

All experiments were performed in a quiet and dimly lit room. In the room, the only source of lighting was the monitor screen, to avoid distractions. The visual stimuli were displayed on a 1920×1080 LCD monitor running at 120 Hz, encompassing a visual angle of approximately 48×30 degrees of visual angle from a distance of 57 cm. Visual stimuli were presented at the center of the screen. The screen background was grey with a luminance of 46.79 cd/m^2^, while each flash was a briefly presented white disc (radius = 10.5 degrees of visual angle, 400 pixels), with a luminance of 98.17 cd/m^2^. The auditory stimuli were presented by means of two speakers positioned behind the screen, centered to correspond to the position on the screen where the visual stimuli were presented (i.e., due to the expected spatial selectivity of serial dependence effects) [1,5,48]. Each auditory event was a pure tone with a frequency (or pitch) of 700 Hz, with 2-ms ramps at the onset and offset to avoid sound distortions. The intensity of the auditory stimuli was set to approximately 65 dB from the position of the participant. These general parameters were the same in all the experiments.

In Exp. 1, the stimuli were uni-sensory streams of either brief visual events (flashes), or brief auditory events (“beeps”). Each event in both the visual and auditory streams lasted for 30 ms, and different events were separated by a jittered interval of 40-70 ms. The events in each stimulus had a presentation frequency range of 10-14 Hz. Each trial included the presentation of three stimuli: “inducer,” “reference,” and “probe,” always in this order. The stimuli could be either visual or auditory, depending on the condition (see below *Procedure* for more information). The inducer included either 7 or 20 flashes or sounds. The reference was constant, and always included 12 flashes or sounds. The probe varied from trial to trial, and could include 6, 8, 12, 16, or 24 flashes or sounds (i.e., one logarithmic unit below and above the reference numerosity, in a Log2 scale). In Exp. 1, in line with previous studies using a similar paradigm [5,40] the serial dependence effect was conceptualized as the influence of the inducer on the perceived numerosity of the subsequent reference stimulus. Note that in this paradigm, even considering the jittering of the stimulus timing, the duration of the sequence is correlated with numerosity, i.e., the higher the numerosity, the longer the duration. Participants could thus alternatively judge duration instead of numerosity. However, considering the task instructions and that judging duration is usually more difficult than judging numerosity [23], this is less likely.

In Exp. 2, we alternated multi-sensory and uni-sensory stimuli in consecutive trials. In this sequence, the multisensory stimuli were considered as inducers, while the subsequent uni-sensory stimuli as references to assess the serial dependence effect. Serial dependence was thus measured as the influence of each component of the multi-sensory stimulus on the subsequent uni-sensory stimulus. The multi-sensory stimuli were composed of an auditory and a visual stream presented simultaneously. The numerosity of each stream was modulated independently, and could be drawn from two different ranges: a low range including numerosities 4, 5, 6, 7, 8, 9, and a high range including 16, 18, 20, 22, 24, 26 events. The two components of the multi-sensory stimuli could thus be congruent, both including either low of high numerosities, or incongruent, with one low and the other high. In the case of incongruent numerosities, the less numerous sequence was centered to the middle point of the more numerous sequence. The uni-sensory stimuli were either visual or auditory streams, and had a numerosity drawn from an intermediate range including 10, 11, 12, 13, 14, and 15 events. Here we used slightly slower sequences to account for the more difficult task performed by participants in Exp. 2 compared to Exp. 1 (numerosity estimation instead of discrimination). Each event in both the auditory and visual streams lasted 30 ms, and different events were separated by a jittered interval of 80-130 ms, resulting in a presentation frequency range of 7.7-12.5 Hz. Before each stimulus, a cue was presented to indicate the sensory modality that participants had to attend. The cue was either a “A” or a “V,” cueing the auditory or the visual modality, respectively. The cue was presented at the center of the screen, replacing the fixation cross.

In Exp. 3 we used the same paradigm as in Exp. 1. The only differences were the addition of electrophysiological (EEG) recording (see below *Electrophysiological recording and analysis*), and a more regular presentation timing of the visual and auditory streams. Namely, instead of jittering the interval between different events in the stream, we instead used 30-ms events separated by fixed interval of 50 ms, corresponding to a presentation frequency of 12.5 Hz. This was done in order to avoid additional noise (i.e., due to variable stimulus timing) when averaging different trials to compute the event-related potentials (ERPs).

### Procedure

***Experiment 1***. In Exp. 1 the participants performed a discrimination task of sequential numerosities. The experiment was divided into two different conditions, performed by the same participants in two separate sessions within the same day. The two conditions were designed to draw attention either to the visual modality only (“uni-sensory task” condition), or to both the visual and auditory modalities (“multi-sensory task” condition). In other words, to trigger either uni-modal attention, or cross-modal attention. In general, the stimulation sequence in each trial started with the presentation of a central fixation cross, which participants were instructed to fixate throughout the task. After a delay of 200 ms, the “inducer” stimulus, including either 7 or 20 events, was presented at the center of the screen. The inducer stimulus was always irrelevant for the task. After an inter-stimulus interval of 400-500 ms, the constant “reference” stimulus was presented (always with a numerosity of 12 events). Finally, after an interval of 500 ms, the “probe” was presented, which was modulated in numerosity in a trial-by-trial fashion (numerosity ranging from 6 to 24 events). At the end of this stimulation sequence, the fixation cross turned red, signaling the participants to provide a response. The participants were instructed to compare the reference and probe stimulus and indicate which one seemed to contain more events (i.e., select the more numerous sequence). The response was provided by pressing a button on a standard keyboard (“2” for the reference, “3” for the probe, as they were the second and third stimulus in each sequence). After providing a response, the next trial started automatically with a delay of 550-650 ms. In the uni-sensory task condition, the inducer could be either visual (a stream of flashes) or auditory (a stream of sounds). The reference and probe were instead always visual streams of flashes, so that participants only needed to explicitly attend the visual modality to perform the task. In the multi-sensory task condition, while the inducer could similarly be either visual or auditory, also the reference and probe could be either visual or auditory, with the constrain that they were never presented both in the same modality. Namely, the sequence could include an auditory reference and a visual probe, or a visual reference and an auditory probe. Doing so, participants were forced to attend both modalities to perform the task. In both conditions there was no time limit to provide a response, but participants were encouraged to respond as fast as they can. In the uni-sensory task condition, participants completed a total of 160 trials (4 blocks of 40 trials), corresponding to 8 repetitions of each combination of inducer numerosity, inducer sensory modality, and probe numerosity (2 x 2 x 5 levels). In the multi-sensory task condition, participants completed a total of 320 trials (8 blocks of 40 trials), corresponding to 8 repetitions of inducer numerosity, inducer sensory modality, reference sensory modality, and probe numerosity (2 x 2 x 2 x 5 levels). No feedback was provided regarding the responses. The procedure of Exp. 1 is depicted in Fig. 1A.

***Experiment 2***. In Exp. 2 the participants performed a numerosity estimation task. In this experiment we aimed at modulating attention to different sensory modalities in a trial-by-trial fashion rather than globally as we did in Exp. 1. To this aim, we alternated multi-sensory (i.e., auditory and visual streams presented simultaneously) and uni-sensory (either visual or auditory streams) stimuli in consecutive trials. Before each stimulus, we presented a cue instructing the participant to attend a specific sensory modality (either visual or auditory; “V” or “A”), so that they explicitly attended and judged only one component of the multisensory stimulus. The experiment was thus structured in pairs of trials. First, a central fixation cross was presented on the screen. Then, after a delay of 600 ms, the cue was presented at the center of the screen, remaining on the screen for the rest of the trial. After 600 ms from the cue onset, a multi-sensory stimulus was presented, with its components including either a low (4-9 events) or a high (16-26 events) numerosity, independently from each other. After the offset of the stimulus, the prompt “Number?” appeared on the screen, and participants were instructed to provide an estimate of the numerosity of the cued component of the stimulus. Participants typed on the keyboard their estimate, and pressed enter to confirm. Only a response between 2 and 40 was accepted, and participants could use the delete button to fix their response in case of a typing error. There was no time limit to provide a response. Even if this was not a speeded task, participants were encouraged to be fast. After providing a response, the fixation cross reappeared for 600 ms and the second trial in a pair started. Again, a cue appeared on the screen, indicating the modality of the subsequent stimulus. Then, after a delay of 600 ms, the uni-sensory stimulus was presented, with a numerosity drawn from an intermediate range compared to the preceding stimulus (10-15 events). The response phase was the same as the multi-sensory stimulus. Participants completed a total of 480 trials (15 blocks of 32 trials, each including 16 pairs of multi-sensory and uni-sensory stimuli). The procedure of Exp. 2 is depicted in Fig. 1B.

***Experiment 3***. The paradigm used in Exp. 3 was the same as Exp. 1 (see Experiment 1 paragraph and Fig. 1A), with only a few differences in the timing of the stimuli to optimize the procedure for EEG recording and analysis. Namely, the inter-stimulus interval between inducer and reference was 600 ms, while the interval between the reference and probe was 700 ms. This was done specifically to have a large enough epoch for the analysis of activity evoked by the reference, before the onset of the subsequent probe. Except for these differences, all other parameters were identical to Exp. 1. The number of trials and repetitions used in Exp. 3 is the same as Exp. 1.

### Behavioral data analysis

In Exp. 1 and Exp. 3, the analysis of behavioral data was aimed at assessing the effect of the task-irrelevant inducer on the perceived numerosity of the subsequent reference stimulus. To do so, we computed the proportion of “probe more numerous” responses at different values of the probe stimulus, as a function of the numerosity of the inducer (7 or 20 events), its modality (visual or auditory), and the modality of the reference (visual or auditory, only in the multi-sensory task condition). Then, we fitted a cumulative Gaussian curve to the proportion of responses, according to the maximum likelihood method [49]. In the fitting procedure, we applied a finger error rate-correction of 2.5%, which proportionally compresses the asymptotic levels of the fit to account for random errors and lapses of attention [50]. From this fit, we computed the point of subjective equality (PSE) as a measure of the perceived numerosity of the reference (i.e., accuracy). Additionally, we computed the just noticeable difference (JND) as the slope of the fit, which in turn was used to compute the Weber’s fraction (WF) to obtain a measure of precision in the task (WF = JND / PSE). To better assess the strength of serial dependence biases, we computed a normalized serial dependence effect index similar to previous studies (e.g., Fornaciai & Park, 2018, 2019a, 2019b), according to the following formula:

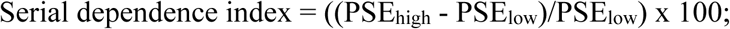

Where the PSE_high_ reflects the PSE obtained with an inducer higher in numerosity compared to the reference (20 events), and PSE_low_ reflects the PSE obtained with an inducer lower in numerosity compared to the reference (7 events). This was done separately for each combination of the inducer and reference sensory modalities in the uni-sensory and multi-sensory task condition. The serial dependence effect indexes were assessed for statistical significance first using one-sample t-tests against a null hypothesis of zero effect. Effects obtained with different inducer and reference sensory modalities were compared using either paired t-tests (in the uni-sensory task) or linear mixed-effect (LME) models (in the multi-sensory task condition). Weber’s fractions across the two tasks were compared with a paired t-test.

In Exp. 2 the analysis involved assessing the effect of the different components of the multi-sensory stimulus (visual and auditory), their combination, and their task-relevance, on the perceived numerosity of the subsequent uni-sensory stimulus. To this aim, we pooled together the trials corresponding to different combinations of the numerosities of the multi-sensory stimulus, the cued modality, and the modality of the uni-sensory stimulus, and computed the average reported numerosity of the uni-sensory stimulus according to these combinations. We then computed a measure of normalized serial dependence effect in percentage similarly to Exp. 1, according to the following formula:

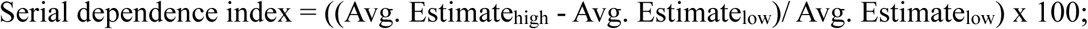

Where the Avg. Estimate_high_ reflects the average numerical estimation of the uni-sensory stimulus obtained where a given component of the preceding multi-sensory stimulus had higher numerosity, and Avg. Estimate_low_ reflects the numerical estimates obtained when the component of the multi-sensory stimulus had lower numerosity.

To evaluate the statistical significance of the effects we used one-sample t-tests against zero. Effects obtained with different combinations of sensory modalities were compared using LME models. Additionally, we also assessed the effects as a function of the congruency of the two components of the multi-sensory stimulus. Namely, when the numerosity of both the auditory and visual component of the stimulus were high vs. low, or when one was high and the other low. In the case of congruent components, we computed the serial dependence effect as described above. In the case of incongruent components, we used a slightly different formula:

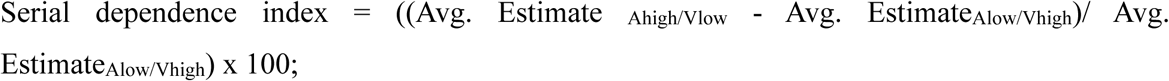

Computing the effect using different combinations of auditory and visual numerosity as we did means that a positive value of the serial dependence effect index reflected the auditory component being dominant, while a negative value reflected the visual component being dominant. The statistical significance of these effects was evaluated using one-sample t-tests against zero, and LME models were used to compare effects as a function of the different sensory modalities involved.

To assess and compare the precision of numerosity estimation in different sensory modalities and at different numerical magnitudes, we computed the Weber’s fraction as the standard deviation of responses at each numerosity level divided by the average estimated number. We then used a LME model to assess potential differences across modalities and numerical magnitudes.

When running multiple t-tests within a condition, multiple comparisons were controlled using a false discovery rate (FDR) procedure, with q = 0.05. In these cases, when reporting the results of the tests we indicate the p-value adjusted with FDR (adj-p).

### Electrophysiological recording and pre-processing

In Exp. 3, during the experimental session we recorded the EEG to address the neural signature of the serial dependence effect within and across vision and audition. The EEG was recorded using the Biosemi ActiveTwo system (2048 Hz sampling rate) and a 128-channel cap based on the 10-20 system layout. To monitor artifacts due to eye movements and blinks, the electro-oculogram (EOG) was measured via an additional electrode attached below the left eye of the subject. The electrode offset values (i.e., the difference in voltage between each electrode and a control electrode) across the channels were usually kept below 20 µV, but occasional values up to 30 µV were tolerated.

Pre-processing of the EEG data was performed offline in Matlab (version R2024b), using the routines provided by the EEGLAB [51] and ERPlab [52] toolboxes. First, EEG signals were re-sampled to a sampling rate of 1000 Hz. Then, we binned each combination of inducer numerosity, inducer sensory modality (and reference sensory modality, only in the multi-sensory task condition) separately. The continuous EEG data was then epoched time-locking the signal to the onset of the reference stimulus, which had the same numerosity in each trial. The epochs spanned from –300 ms to 1500 ms around the stimulus onset. The pre-stimulus interval (−300:0 ms) was used for baseline correction. The EEG signal was band-pass filtered with cut-offs at 0.1 and 40 Hz. To reduce artefactual activity in the data, we used an independent component analysis (ICA), aimed at removing identifiable artifacts such as eye movements and blinks. We additionally employed a step-like artifact rejection procedure (amplitude threshold = 100 μV, window = 400 ms, step = 20 ms) to further remove any remaining large artifact from the signal, leading to the exclusion of 0.63% ± 0.81% of the trials, on average (± SD). Finally, the event-related potentials (ERPs) were computed by averaging EEG epochs within each bin. Doing so, we obtained a measure of the average brain responses to the reference stimulus as a function of the preceding inducer, and as a function of the different sensory modalities of the stimuli (only the inducer modality in the uni-sensory task, and the modality of inducer and reference in the multi-sensory task). ERPs were further low-pass filtered with a cut-off at 30 Hz. All statistical tests described in the next paragraphs (t-tests, linear mixed-effect model) were performed considering a series of small time windows (50 ms with a step of 5 ms) throughout the epoch in a sliding-window average fashion, to increase the signal-to-noise ratio. The ERP contrasts plotted in Fig. 5D, Fig. 6F, and Fig. 6H are smoothed with a with a sliding-window average with a width of 50 ms and a step of 5 ms to reflect the time windows used in the statistical tests. When reporting the results of tests, we list the center of the first and last latency window of the significant cluster. Similarly, the indications of significant latency windows in the figures (thick lines) start and end at the center of the first and last significant window in the cluster.

The channels of interest used for visualization of the ERPs and for data analysis were selected by examining the topographic distribution of activity evoked by the reference stimulus. Since serial dependence is defined in our paradigm as the effect of the inducer on the perceived numerosity of the reference, the rationale for doing so was to select the channels best capturing the responses to the reference stimulus itself. We thus selected a group of channels showing the peak of activity in response to the reference stimulus, evaluated within the reference presentation epoch. In both the case of the visual and auditory reference stimuli, we accordingly selected a set of 12 central occipital and occipito-parietal channels, including I1, Iz, I2, OI1, OIz, OI2, O1, Oz, O2, PO1, POz, PO2. These channels are also consistent with previous studies addressing the neural signature of numerosity perception [53–57] and serial dependence in numerosity perception [37,38].

The significance of the effect of the inducer numerosity on ERPs was assessed by comparing the ERP amplitude in the case of a low versus a high inducer, using a series of paired t-tests. To control for multiple comparisons, we used cluster-based non-parametric tests. Namely, first, we set a cluster threshold of at least three consecutive significant time-windows (i.e., only clusters of at least three consecutive significant tests were considered in data analysis). Then, for each cluster of consecutive significant time windows, we ran a series of permutation tests. Namely, at each time window within the cluster, we selected two random halves of the datasets to be compared, and mixed them. The paired t-test was performed on these shuffled datasets. This procedure was repeated 10,000 times, selecting different subsets of the data and repeating the test. Finally, for each repetition, we computed the total t-value across the simulated cluster, and compared it to the minimum t-value obtained in the real cluster (to be more conservative) multiplied by the number of time windows. The proportion of times in which the total t-value of the simulated cluster was equal or higher than the actual cluster was taken as the p-value of the test.

To perform further analyses, we also computed a measure of ERP contrast, which reflects the difference in ERP amplitude between the brain responses to the reference preceded by a high versus a low inducer. First, the contrast amplitude in different conditions (e.g., visual vs. auditory inducer), reflecting the differences in brain responses to the reference as a function of the inducer, was tested with paired t-tests. In the uni-sensory task condition, we compared the contrast relative to the visual versus the auditory inducer. In the multi-sensory task condition, we performed similar comparisons, but separately for the visual and auditory reference, since visual and auditory stimuli likely result in different patterns of activation independently from the preceding inducer. This index was also computed to further assess the relationship between the ERP signature of the effect of the inducer, and the behavioral serial dependence effect. Similarly to the ERP contrast, the serial dependence effect in this case was defined as the difference in PSE corresponding to a high versus a low inducer. To do so, we used a series of LME models within each time window throughout the reference stimulus epoch, adding the behavioral effect as the dependent variable, the ERP contrast as the fixed effect, and the subjects as the random effect. For both the paired t-tests comparing the contrast amplitudes and the LME models, multiple comparisons were controlled with cluster-based non-parametric tests, according to the same procedure described above for the comparison of ERPs. In the case of the LME models, the only difference in the procedure was that we shuffled both the design matrix and the data entered into the model prior to running the simulated test.

### Data availability

All the data generated during the experiments described in this manuscript is publicly available on Open Science Framework, at this link: https://osf.io/9y3e7/.

## Authors contributions

IT, MF, and OC designed the study; IT and MF programmed the stimuli and procedure; IT, MF, and SB collected the data; IT, MF, and SB analysed the data; IT, MF, SB, and OC wrote the initial draft of the manuscript; IT, MF, and OC revised the manuscript.

